# Metabolic dissimilarity determines the establishment of cross-feeding interactions in bacteria

**DOI:** 10.1101/2020.10.09.333336

**Authors:** Samir Giri, Leonardo Oña, Silvio Waschina, Shraddha Shitut, Ghada Yousif, Christoph Kaleta, Christian Kost

## Abstract

The exchange of metabolites among different bacterial genotypes profoundly impacts the structure and function of microbial communities. However, the factors governing the establishment of these cross-feeding interactions remain poorly understood. While shared physiological features may facilitate interactions among more closely related individuals, a lower relatedness should reduce competition and thus increase the potential for synergistic interactions. Here we investigate how the relationship between a metabolite donor and recipient affects the propensity of strains to engage in unidirectional cross-feeding interactions. For this, we performed pairwise cocultivation experiments between four auxotrophic recipients and 25 species of potential amino acid donors. Auxotrophic recipients grew in the vast majority of pairs tested (78%), suggesting metabolic cross-feeding interactions are readily established. Strikingly, both the phylogenetic distance between donor and recipient and the dissimilarity of their metabolic networks were positively associated with the growth of auxotrophic recipients. Analysing the co-growth of species from a gut microbial community *in-silico* also revealed that recipient genotypes benefitted more from interacting with metabolically dissimilar partners, thus corroborating the empirical results. Together, our work identifies the metabolic dissimilarity between bacterial genotypes as key factor determining the establishment of metabolic cross-feeding interactions in microbial communities.

**Highlights:**

- The exchange of essential metabolites is common in microbial communities
- Metabolic cross-feeding interactions readily establish between auxotrophic and prototrophic bacterial strains
- Both the phylogenetic and the metabolic dissimilarity between donors and recipients determines the successful establishment of metabolic cross-feeding interactions

## Introduction

Microorganisms are ubiquitous on our planet and are key for driving pivotal ecosystem processes [1–3]. They contribute significantly to the flow of elements in global biogeochemical cycles [3, 4] and are also crucial for determining the fitness of plants [5, 6] and animals [7, 8], including humans [9, 10]. These vital functions are provided by complex communities that frequently consist of hundreds or even thousands of metabolically diverse strains and species [11, 12]. However, the rules that determine the assembly, function, and evolution of these microbial communities remain poorly understood. Understanding the underlying governing principles is central to microbial ecology and crucial for designing microbial consortia for biotechnological [13] or medical applications [14, 15].

In recent years, both empirical and theoretical work has increasingly suggested that the exchange of essential metabolites among different bacterial genotypes is a crucial process that can significantly affect growth [16, 17], composition [18], and the structure of microbial communities [19]. In these cases, one bacterial genotype releases a molecule into the extracellular environment, which can then be used by other cells in the local vicinity. The released substances frequently include building block metabolites such as amino acids [20, 21], vitamins [22, 23], or nucleotides [24], as well as degradation products of complex polymers [19, 25]. Even though these compounds represent valuable nutritional resources, they are released as unavoidable byproducts of bacterial physiology [26, 27] and metabolism [28] or due to leakage through the bacterial membrane [29, 30]. Consequently, the released compounds create a pool of resources that can benefit both conspecifics and members of other species in the local vicinity [31–34]. The beneficiaries include genotypes that opportunistically take advantage of these metabolites and strains, whose survival essentially depends on an external supply with the corresponding metabolite. Due to a mutation in their genome, these so-called *auxotrophic* genotypes are unable to autonomously synthesise vital nutrients such as amino acids, vitamins, or nucleotides. By utilising metabolites that are produced by another cell, a unidirectional cross-feeding interaction is established. Auxotrophic mutants that use compounds released by others gain a significant fitness advantage over prototrophic cells that produce the required metabolites by themselves [35]. Due to the strong fitness benefits that can result for auxotrophic genotypes, this type of cross-feeding interaction is prevalent in all kinds of microbial ecosystems, including soil [36], fermented food [21], aquatic environments [37, 38], as well as host-associated microbiota [7, 39]. Despite the ubiquity of unidirectional cross-feeding interactions in nature, the factors determining their establishment remain poorly understood [40–43]. In particular, it is unclear how the relationship between the metabolite donor and the auxotrophic recipient affects the likelihood that a cross-feeding interaction is successfully established. Two possibilities are conceivable.

First, phylogenetically more closely related individuals are likely to share physiological features that favour the establishment of cross-feeding interactions relative to strains lacking these attributes. For example, an efficient transfer of metabolites from one cell to another commonly depends on close physical contact between donor and recipient [44]. The attachment to other cells is generally mediated by surface factors (e.g. adhesive proteins) or exopolymers that, in some cases, operate more effectively between cells sharing similar surface structures [45]. Moreover, some species form intercellular nanotubes to exchange metabolites between cells [44, 46], which in turn might require an increased structural similarity between cells for efficient transport to operate. Another context, in which the phylogenetic relatedness between donor and recipient could determine the establishment of a unidirectional cross-feeding interaction, is when both partners can communicate with each other. Certain signals that are involved in chemical communication (i.e. quorum sensing) between cells are more readily perceived by more closely related bacterial strains than more distantly related individuals [47]. Consequently, quorum sensing between more similar genotypes is also more likely to regulate processes such as cell-cell adhesion [16] or the establishment of metabolic cross-feeding interactions [48]. In the following, we refer to this possibility as the *similarity hypothesis.*

Second, unidirectional cross-feeding interactions might favour more distantly related donor-recipient pairs over interactions among close relatives. Two closely related bacterial cells are more likely to share ecological preferences such as habitat or resources utilised than two phylogenetically different bacterial taxa [41, 49, 50]. Moreover, two phylogenetically, more closely related cells will tend to have a more similar metabolic network than two distantly related cells [33, 51, 52]. Consequently, both the biosynthetic cost to produce a given metabolite and its nutritional value are more likely to differ in heterospecific pairs than amongst members of the same species [53, 54]. If these differences also translate into an enhanced growth of the auxotrophic recipient, a positive correlation between the growth of the auxotroph and the phylogenetic and/ or metabolic distance to the donor cell would be observed. In the following, we refer to this alternative possibility as the *dissimilarity hypothesis.*

Here we aim to distinguish between these two hypotheses to better understand the factors governing the establishment of this ecologically very important interaction. To achieve this goal, we use unidirectional cross-feeding interactions as a model. Synthetically assembling pairs consisting of an auxotrophic recipient and a prototrophic amino acid donor of the same or a different species ensured that both interaction partners do not share a coevolutionary history. In this way, all results will represent the situation of a naïve encounter between both interaction partners and only mirror effects resulting from the phylogenetic relatedness and metabolic dissimilarity between partners. Using this synthetic ecological approach, we systematically determined whether and how the phylogenetic or metabolic distance between auxotrophic recipients and prototrophic amino acid donors affects cross-feeding in pairwise bacterial consortia.

Our results show that in the vast majority of cases tested, unidirectional cross-feeding interactions successfully established between a prototrophic donor and an auxotrophic recipient. Strikingly, recipients’ growth was positively associated with both the phylogenetic and metabolic distance between donor and recipient. This pattern could partly be explained by the difference in the amino acid profiles produced by donors. Finally, analysing the co-growth of species from a gut microbial community *in-silico* revealed that recipient genotypes benefitted more from interacting with metabolically dissimilar partners, thus corroborating the empirical results. Together, our work identifies the metabolic dissimilarity between donor and recipient genotypes as a critical parameter determining the establishment of unidirectional cross-feeding interactions in microbial communities.

## Results

### Auxotrophic recipients commonly benefit from the presence of prototrophic donor cells

To determine the probability with which unidirectional cross-feeding interactions emerge between an auxotrophic recipient and a prototrophic donor genotype, pairwise coculture experiments were performed (Fig. 1A). For this, 25 strains that belonged to 21 different bacterial species were used as potential amino acid donors (Figs. 1B and S1A). Donor strains were selected such that they represented different bacterial taxa and were able to synthesise all nutrients they required for growth in a minimal medium (metabolic autonomy, prototrophy). These potential amino acid donors were individually cocultured together with each one of four auxotrophic recipients that belonged to one of two bacterial species (i.e. *Escherichia coli* and *Acinetobacter baylyi)* and were auxotrophic for either histidine (Δ*hisD*) or tryptophan (Δ*trpB*) (Fig. S1B).

**Fig. 1.**
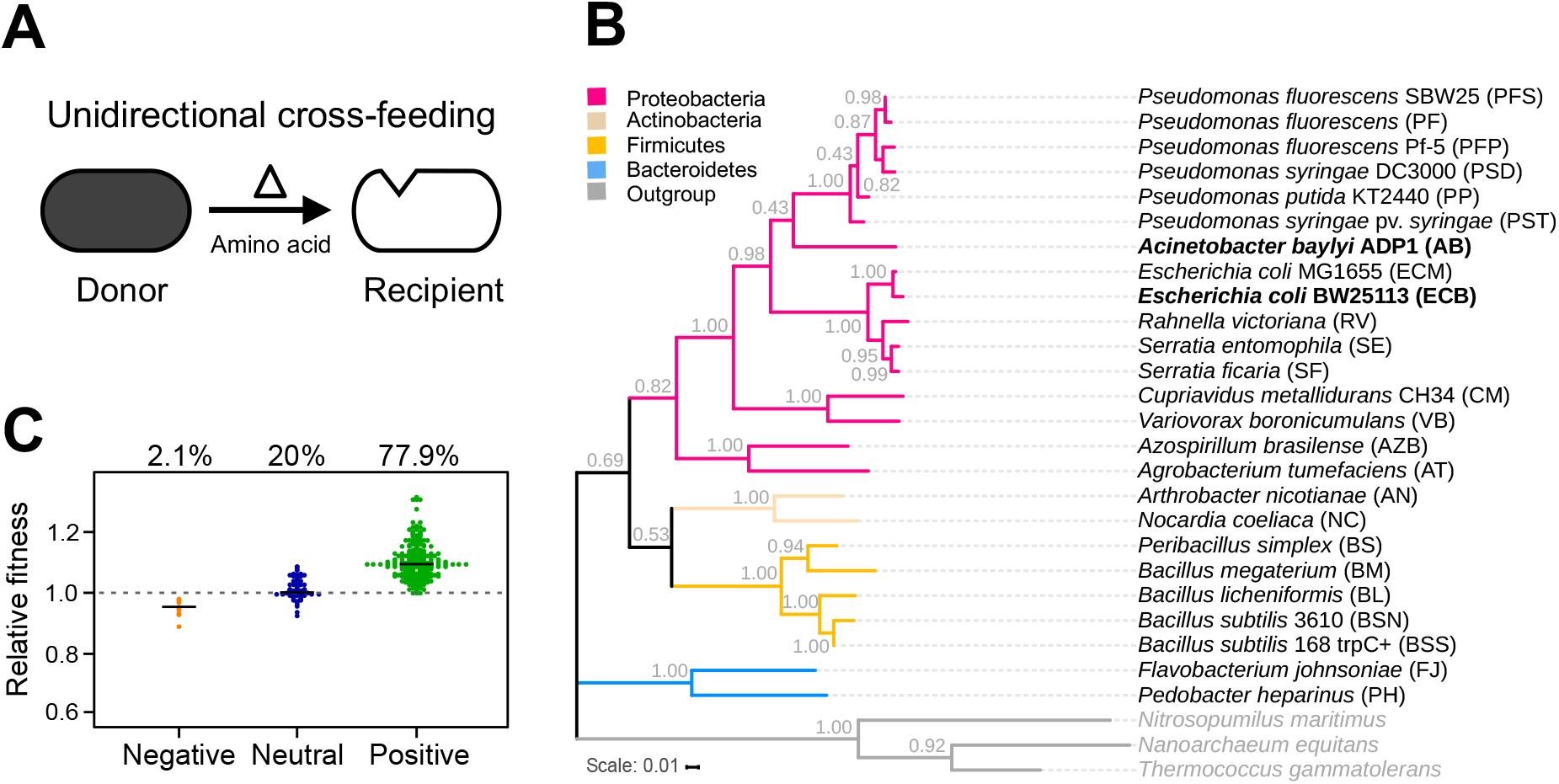
Unidirectional cross-feeding between prototrophic donor cells and amino acid auxotrophic recipients is common. **(A)** Overview over the experimental system used. Metabolically autonomous donor genotypes (dark cell) were cocultivated together with an auxotrophic recipient that was unable to produce either histidine or tryptophan (white cell). Growth of auxotrophs signifies the successful establishment of a unidirectional cross-feeding interaction, in which the focal amino acid (e.g. histidine (△)) is exchanged between donor to recipient cells. **(B)** Phylogenetic tree of bacterial species (donors and recipients) used in this study. Different colors indicate different phyla. The tree was constructed based on the 16S rRNA gene. Recipient strains used in this study are highlighted in bold. Branch node numbers represent bootstrap support values. **(C)** Growth of auxotrophic recipients in pairwise coculture with different donor genotypes. *Escherichia coli* and *Acinetobacter* baylyi, each either auxotrophic for histidine (Δ*hisD)* or tryptophan (Δ*trpB),* were used as amino acid recipients. The relative fitness of receivers, when grown in coculture with one of 25 donors, is plotted relative to their growth in monoculture in the absence of the focal amino acid (dashed line). CFU was calculated 24 h post-inoculation. Interactions in cocultures were classified as negative (n = 8), neutral (n = 76), and positive (n = 296), based on the statistical difference between the growth of auxotrophs in monoculture and coculture (FDR-corrected paired t-test: P ≤ 0.05).

To test if the selected donor strains can support the growth of auxotrophic recipients, the abovementioned strains were systematically cocultured in all possible pairwise combinations (initial ratio: 1:1). Subsequently, the growth of the recipient strains in coculture was quantified at the onset (0 h) and after 24 h and compared to the growth, the same strain achieved in a monoculture over the same period in the absence of externally supplied amino acids. In this experiment, the donor’s presence affected the recipient’s growth either positively, negatively, or in a neutral way. Only 2% of the tested cases showed a growth reduction, and in 20% of interactions, auxotrophs did not respond at all to the presence of a donor cell (Fig. 1C). In contrast, in the vast majority of cocultures tested (i.e., 78%), the growth of auxotrophic cells was significantly enhanced in the presence of donor cells as compared to their growth in monocultures, suggesting that unidirectional cross-feeding interactions can readily establish (Fig. 1C).

### Recipient growth depends on amino acid production of donor genotypes

The main factor causing the growth of auxotrophs in the coculture experiments was likely the amount and identity of metabolites that donor cells released into the extracellular environment (i.e., the exo-metabolome) [26]. To test if amino acid production of donors could explain the observed recipient growth, the supernatant of monocultures of all 25 donor strains was collected during exponential growth. Subjecting the cell-free supernatant of these cultures to LC-MS/MS analysis revealed that all tested genotypes secreted amino acids in varying amounts (Figs. 2A and S2). In this experiment, donors are not expected to specifically produce the amino acid that the cocultured auxotroph requires for growth. Moreover, bacteria usually use generic transporters to import chemically similar amino acids [55–57]. Thus, auxotrophic recipients may benefit not only from the one amino acid they require for growth but potentially also from utilising other amino acids that are produced by the donor. To quantitatively determine whether the released amino acids could explain the observed growth of recipients, the cell-free supernatant of donor cultures (replenished with fresh nutrients, see methods) was supplied to monocultures of auxotrophic cells, and the resulting growth over 24 h was quantified. In addition, the chemical composition of the supernatants used was determined via LC-MS/MS. As expected, the growth of auxotrophic recipients was positively associated with the concentration of the amino acid the corresponding auxotroph required for growth (Fig. 2B). Interestingly, however, was the observation that recipient growth also correlated positively with the total amount of amino acids present in the donor supernatant (Fig. 2B). Together, these results show that auxotrophic recipients not only use the amino acids they cannot produce autonomously but also take advantage of the other amino acids produced by donor cells.

**Fig. 2.**
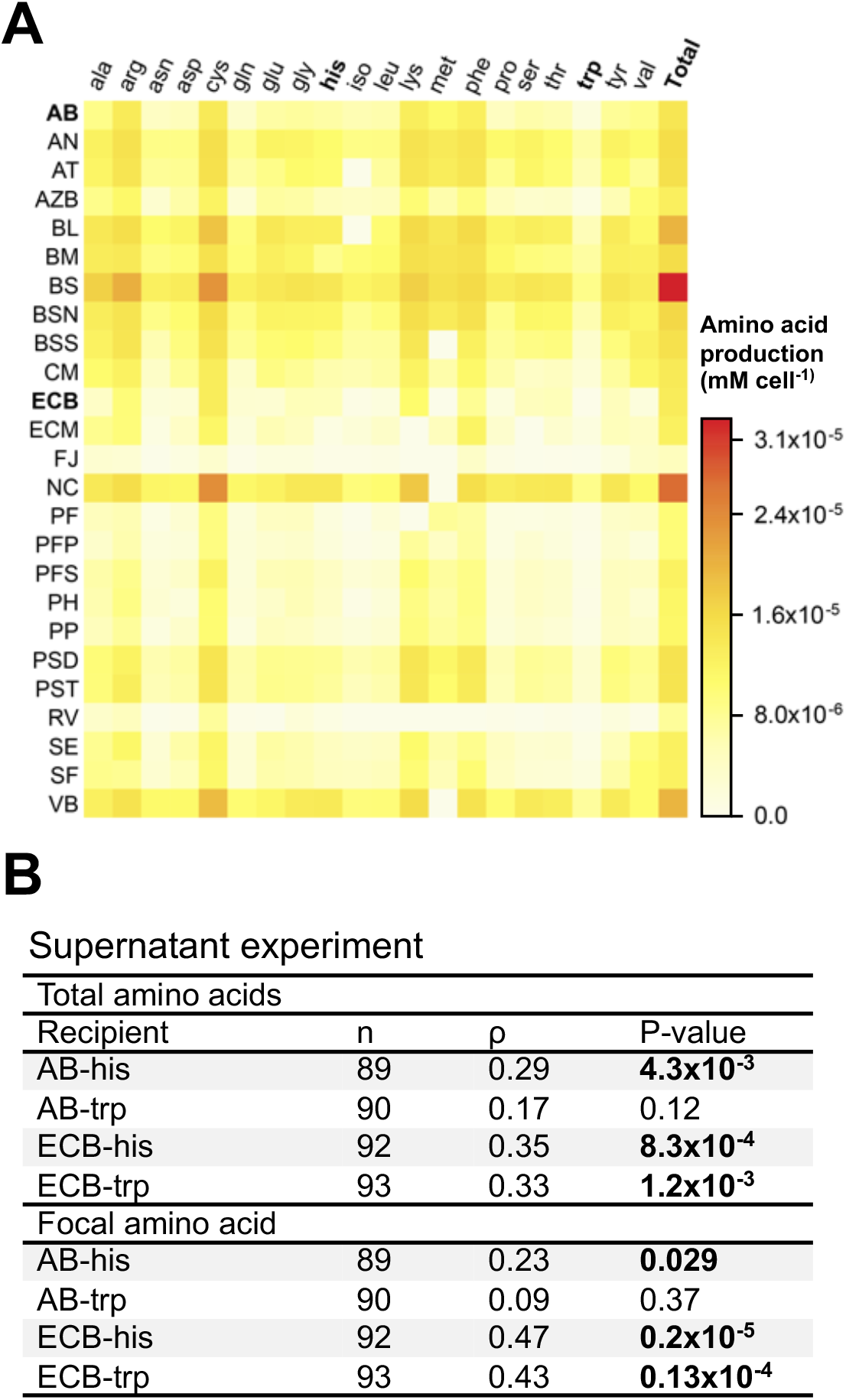
Total amino acid production of different donors can predict unidirectional cross-feeding. **(A)** Heatmap of amino acids released by different donor strains. Amount of amino acid (mM per cell) produced by 25 donor strains (for abbreviations see Fig. 1B) is shown (Y-axis). Cell-free supernatants of exponentially growing cultures were analysed via LC-MS/MS. Colours indicate different amino acid concentrations (legend) and the total amino acid produced by the donor. The two focal amino acids used in the experiments are highlighted in bold letters (i.e. his and trp) **(B)** Overview over the statistical relationships between the total amount of amino acids (upper part) or the focal amino acid (lower part) in supernatants of donor cultures and the growth of the corresponding auxotrophic recipients. Results of spearman rank correlations (⍴) are shown.

### Recipient growth correlates positively with amino acid profile dissimilarity

To distinguish between the two main hypotheses, we asked whether the difference in the amino acid profile (i.e., the collection of amino acids secreted by the donor cell) produced by a closely and distantly related donor strain could explain the growth auxotrophs achieved in the coculture experiment. To test this, we calculated the Euclidean distance between the amino acid profiles of all 25 donor strains. Comparing the statistical relationship between the normalised growth of auxotrophs in coculture with the Euclidean distance in the amino acid profiles of closely and distantly related donor genotypes revealed a significantly positive relationship between both parameters in all cases (Fig. 3A and Table 1). In other words, auxotrophs grew better in coculture with a donor, which contained an amino acid mixture whose composition was different from the one a conspecific cell would have produced. Thus, these results support the dissimilarity hypothesis.

**Fig. 3.**
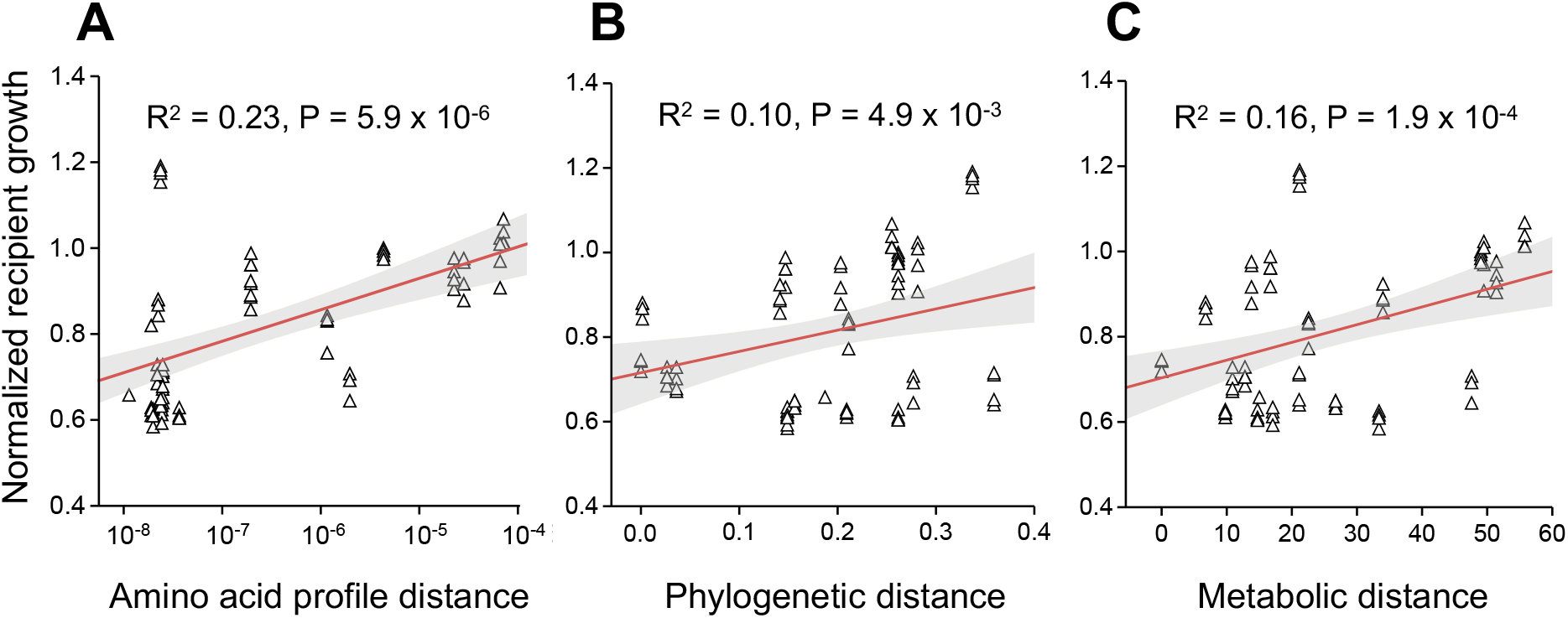
Cross-feeding increases with an increasing dissimilarity to donor cells. Shown is the net growth of the *E. coli* recipient auxotrophic for histidine (Δ*hisD*, △) as a function of **(A)** the amino acid profile distance, **(B)** the phylogenetic distance, and **(C)** the genome-based metabolic distance between donor and recipient. Red lines are fitted linear regressions, and grey area indicates the 95% confidence interval. Each triangle (△) represents a replicate and the sample size is 80 in all cases. Growth of recipient is displayed as a logarithm of the difference in the number of CFUs between 0 h and 24 h and was normalised per number of donor cells.

**Table 1.** Amino acid profile distance (AAD), phylogenetic distance (PD), and metabolic distance (MD) are positively associated with recipient growth. Results of statistical regressions are shown.

| Recipient | n | AAD |  | PD |  | MD |  |
| --- | --- | --- | --- | --- | --- | --- | --- |
|  |  | R<sup>2</sup> | P-value | R<sup>2</sup> | P-value | R<sup>2</sup> | P-value |
| AB-his | 87 | 0.16 | **8.0x10<sup>-5</sup>** | 0.45 | **1.61x10<sup>-12</sup>** | 0.26 | **3.82x10<sup>-7</sup>** |
| AB-trp | 100 | 0.07 | **5.5x10<sup>-3</sup>** | 0.11 | **7.92x10<sup>-4</sup>** | 0.19 | **5.59x10<sup>-6</sup>** |
| ECB-his | 80 | 0.23 | **5.9x10<sup>-6</sup>** | 0.10 | **4.95x10<sup>-3</sup>** | 0.16 | **1.96x10<sup>-4</sup>** |
| ECB-trp | 91 | 0.26 | **2.1x10<sup>-7</sup>** | 0.12 | **6.40x10<sup>-4</sup>** | 0.17 | **3.80x10<sup>-5</sup>** |

### Growth of recipients scales positively with the phylogenetic and metabolic distance to donor cells

Next, we asked whether two phylogenetically close genotypes are more likely to engage in a unidirectional cross-feeding interaction than two more distantly related genotypes. To test this, we re-analysed the results of the coculture experiment by focusing on the phylogenetic relatedness between donor and recipient genotypes. In this context, only those cocultures were considered, in which auxotrophs showed detectable growth. These analyses revealed a positive association between the recipients’ growth and its phylogenetic distance to donor cells (Fig. 3B and Table 1).

However, given that previous analyses suggested differences in the amino acid profiles could predict the growth of auxotrophic recipients (Fig. 3A and Table 1), we reasoned that the phylogenetic distance might only approximate the difference in the strains’ metabolic networks. To verify this, we compared the genome-scale metabolic networks of all donor genotypes. A metabolic similarity matrix between donor and recipient strains was calculated by identifying similarities and differences in both partners’ biosynthetic pathways. Correlating the resulting data with the growth of auxotrophic recipients in coculture revealed a positive association between the metabolic distance and recipient growth (Fig. 3C and Table 1). Together, these results provide additional support for the hypothesis that cross-feeding interactions are more likely to establish between two more dissimilar genotypes.

### All three distance measures alone can explain recipient growth

Having observed a significant positive correlation of recipient growth with each of the three-distance metrics analysed (i.e., amino acid profile distance (AAD), phylogenetic distance (PD), and metabolic distance (MD), Table 1), we asked whether these factors alone were sufficient to predict the growth of auxotrophic recipients. This question was addressed by replotting the data of the performed coculture experiments in three-dimensional graphs that display the growth of a given auxotroph depending on two of the three measures quantified. Fitting a 2D plane into the resulting graphs indicated that increasing each of the three measures also increased recipients’ growth (Fig. 4 and S3). Thus, these graphs suggested that the three explanatory variables are likely correlated with each other. To subject this conjecture to a formal statistical test, we repeated the regression analyses to examine whether MD or AAD was significantly associated with auxotrophs’ growth in coculture, when the first predictor variable PD was already included (Table S1). In all cases, the growth of *E. coli* recipients remained positively associated with metabolic distance as well as the distance of the amino acid production profile (Table S1). However, the tryptophan auxotroph of *A. baylyi* (Δ*trpB*) showed only marginally significant effects, while the pattern no longer held for the histidine auxotroph (Δ*hisD*) (Table S1).

**Fig. 4.**
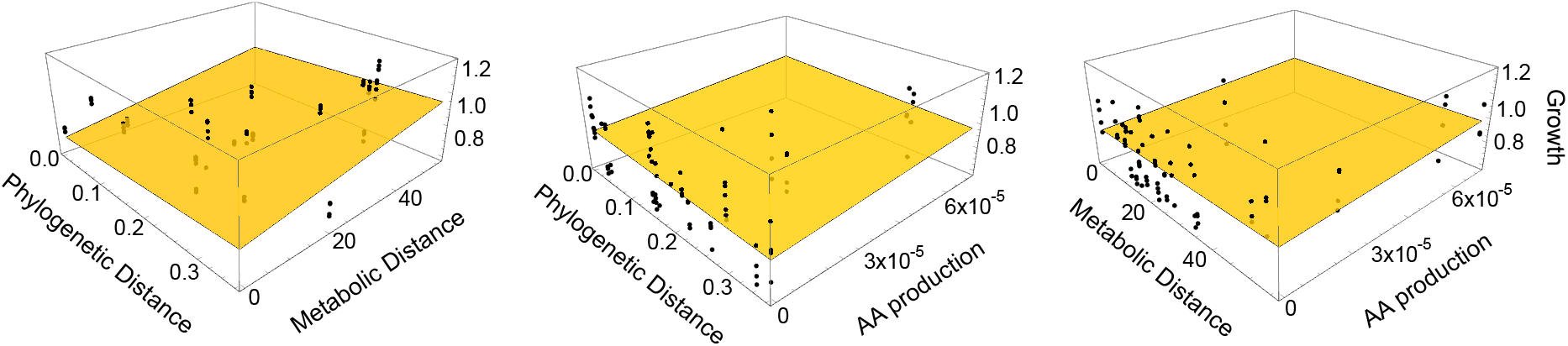
Multiple distance measures interactively explain recipient growth. The plane depicts the linear regression between the growth of the histidine auxotrophic *E. coli* recipient (Δ*hisD*) and the phylogenetic distance, the metabolic distance, and the amino acid profile distance between donor and recipient. Data points above the plane are shown in black. See also Fig. S3 for other comparisons.

Next, we asked whether the amount of amino acid produced by donors was sufficient to explain the growth observed in auxotrophic recipients. One possibility could have been that the positive association between the three distances measures (i.e., amino acid profile distance (AAD)) phylogenetic distance (PD), and metabolic distance (MD)) showed with recipient growth was, in fact, only due to a positive correlation of these parameters with the amount of amino acids produced by donor genotypes. To control this, we first calculated a linear regression for both AAD, PD, and MD with the total amount of amino acids (TAA) or the amount of the focal amino acid (FAA) produced as the independent variable. The residuals (i.e., the variation not explained by TAA or FAA) were then used as independent variables in the regressions. In almost all cases, recipients’ growth remained significantly positively associated with the three distance measures (Table S2 and S3). Together, the set of analyses performed demonstrates that the three different measures analysed (i.e., AAD, PD, and MD) can individually (in the case of *E. coli*) or in combination (both species) explain the cross-feeding between prototrophic donors and auxotrophic recipients, thus corroborating the dissimilarity hypothesis.

### *In-silico* model confirms that the metabolic dissimilarity between species enhances cross-feeding

To verify whether the patterns observed in laboratory-based coculture experiments also applied to natural microbial communities, *in-silico* modelling was used to simulate the co-growth of different bacterial species that co-occur in the human gastrointestinal tract. Specifically, all 334,153 pairwise combinations of 818 bacteria commonly found in this environment were considered. The *in-silico* simulations indicated that the relationship between metabolic as well as phylogenetic distance and the frequency of pairs, for which at least one of the organisms gains a growth advantage from a metabolic interaction in coculture, follows a saturation curve (Fig. 5 and Table S4). This finding shows that bacteria residing within the human gut are more likely to engage in cross-feeding interactions with metabolically more dissimilar species. Taken together, the set of computational analyses performed here is in line with the experimental data shown above: both datasets reveal that metabolic cross-feeding interactions are more likely to establish between two metabolically more dissimilar partners.

**Fig. 5.**
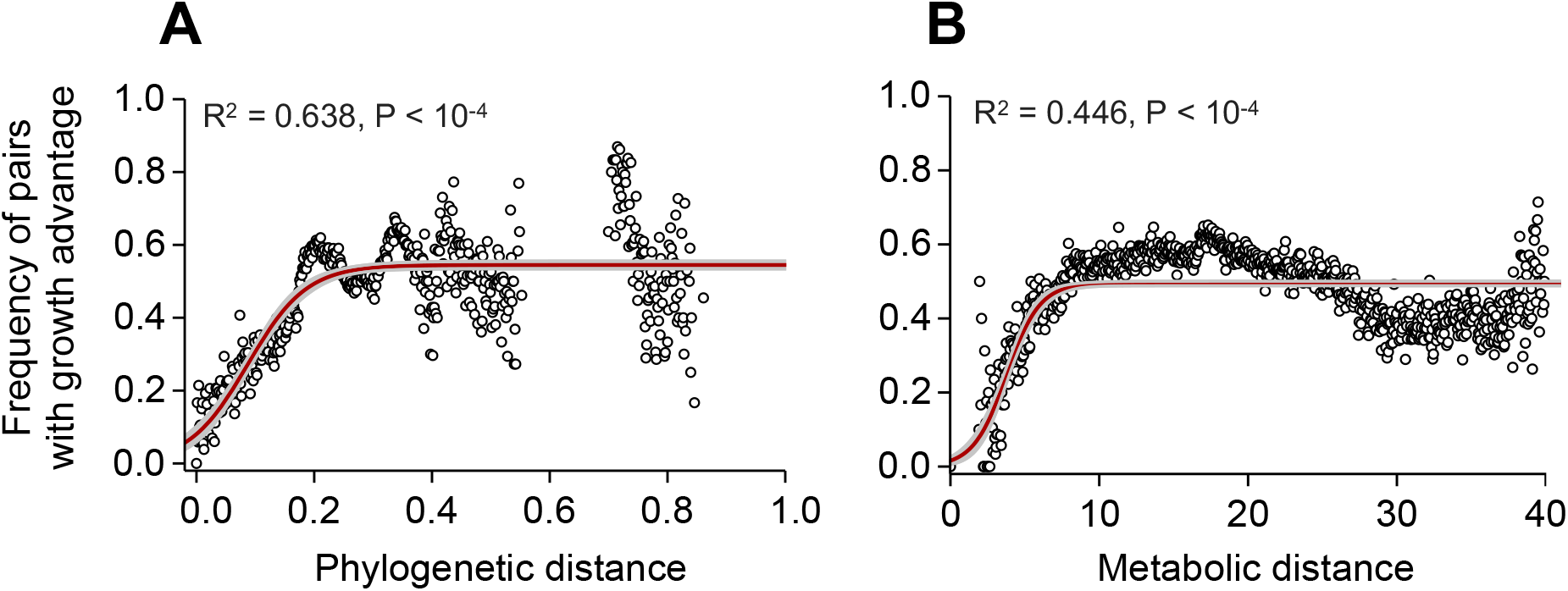
Metabolic simulations of gut bacterial cocultures predict a growth advantage that increases with increasing metabolic and phylogenetic distance. Shown are the results of an *in-silico* flux-balance-analysis of paired models analysing 334,153 combinations of 818 bacterial species residing in the human gut. A pair of species is considered as a pair with growth advantage, if at least one of the organisms is predicted to grow better in cocultures as compared to the predicted growth rate in monoculture. The frequency of pairs with growth advantage is estimated as a function of the (**A**) phylogenetic and (**B**) metabolic distance by defining 1,000 buckets of uniform distance widths spanning the range from 0 to the largest distance. Bucket values are only shown and included in logistic curve fitting if the bucket included at least 10 species pairs. The red line shows the optimal fit of a logistic model to the data with the SE-interval as grey ribbon (see also Table S4).

## Discussion

Metabolic cross-feeding interactions among different microbial species are ubiquitous and play critical roles in determining the structure and function of microbial communities [16, 58, 59]. However, the rules that govern their establishment remain poorly understood. Here we identify the metabolic dissimilarity between donor and recipient genotype as a major determinant for the establishment of obligate, unidirectional cross-feeding interactions between two bacterial strains. In systematic coculture experiments between a prototrophic amino acid donor and an auxotrophic amino acid recipient, we show that growth of auxotrophic recipients in coculture was positively associated with (i) the compositional difference in the amino acid mixtures various donor produced (Fig. 3A), (ii) their phylogenetic distance (Fig. 3B), as well as (iii) the difference in their metabolic networks (i.e. their metabolic distance) (Fig. 3C). Furthermore, *in-silico* simulations of the co-growth of species from a gut microbial community corroborated that the propensity of cross-feeding interactions to establish increased when both interacting partners were metabolically more dissimilar (Fig 5).

In our study, we manipulated the relatedness between donor and recipient genotypes. A high phylogenetic relatedness between two genotypes (donor and recipient) in coculture means that they can perform similar metabolic reactions and are more likely to be characterised by overlapping growth requirements [33, 43]. Consequently, both the nutritional value of a given molecule and the biosynthetic cost to produce it is alike [53, 54, 60]. In contrast, two phylogenetically distant strains likely differ in their metabolic capabilities and requirements. Thus, two more closely related strains are likely to compete for environmentally available nutrients and provide an increased potential for a difference in the cost-to-benefit-ratio than two distant relatives [43, 61, 62]. This statistical relationship can explain why in our coculture experiments, both the phylogenetic and metabolic distance were positively associated with the growth of cocultured auxotrophs. Thus, our results support the dissimilarity hypothesis to explain the establishment of unidirectional cross-feeding interactions. Our findings are in line with previous studies that analysed the effect of the phylogenetic relatedness and metabolic dissimilarity on antagonistic interactions between two different genotypes. These studies found that bacteria mainly inhibit metabolically more similar and related species [41, 63]. Even though the focal biological process differs drastically between our (metabolic cross-feeding) and these other studies (antagonistic interactions), the main finding is conceptually equivalent: genotypes are more likely to compete against closer relatives, yet support the growth of more dissimilar strains – either by enhancing their growth (Fig. 3) or inhibiting them less [41, 63].

In our experiments, we took advantage of synthetically assembled pairwise interactions between different bacterial genotypes to assess how the similarity between interacting partners affects the cross-feeding of metabolites. Even though this approach is limited by the number of pairwise comparisons that can be analysed in one experiment, the obtained results provide a very clear answer to the focal question. First, the selected donor strains covered a broad range of taxonomic diversity in bacteria (Fig. 1B). Thus, the spectrum of ecological interactions analysed here likely reflects the range of interactions a given bacterial genotype would typically experience in a natural microbial community. Second, by deliberately choosing strains that lack a previous coevolutionary history, any result observed can be attributed to the focal, experimentally-controlled parameter (e.g., phylogenetic or metabolic distance). In this way, confounding effects like an evolved preference for a particular genotype can be ruled out. Finally, we analysed bacterial consortia in a well-mixed, spatially unstructured environment, in which the exchanged metabolites are transferred between cells via diffusion through the extracellular environment. Such a set-up minimises factors that would be amplified in a spatially structured environment, such as a local competition for nutrients or the release of metabolic waste products that inhibit the growth of other cells in the local vicinity. Thus, the experimental approach chosen circumvents the challenges of manipulating and detecting metabolite exchange in natural environments and instead capitalises on analysing experimentally arranged and carefully controlled coculture experiments.

The guiding principle discovered in this study is most likely relevant for ecological interactions outside the realm of microbial communities. Mutualistic interactions, in which two partners reciprocally exchange essential metabolites or services, usually involve two or more completely unrelated species [61, 64–66]. In contrast, cooperative interactions among closely related individuals usually rely on the uni- or bidirectional exchange of the same commodity or service [61]. Thus, two more dissimilar individuals have an increased potential to engage in a synergistic interaction than two more similar individuals may be a universal rule that guides the establishment of mutualistic interactions in general [43].

Our results highlight the utility of using synthetic, laboratory-based model systems to understand the fundamental principles of microbial ecology. In this study, we demonstrated that simple assembly rules likely determine the establishment of interactions in natural microbial communities. These insights enrich our understanding of the complex relationships among bacteria in their natural environment and will help to rationally design and modify them for biotechnological or medical applications.

## Material and methods

### Bacterial strains and their construction

Twenty-five bacterial wild type strains were used as potential amino acid donors (Supplemental Table S5). *Escherichia coli* BW25113 and *Acinetobacter baylyi* ADP1 were used as parental strains, from which mutants that are auxotrophic for histidine (Δ*hisD*) or tryptophan (Δ*trpB*) were generated. The gene to be deleted to create the corresponding auxotrophy was identified using the KEGG [67] and the EcoCyc [68] database. For *E. coli*, deletion alleles were transferred from existing single-gene deletion mutants (i.e. the Keio collection, [69]) into *E. coli* BW25113 using phage P1-mediated transduction [70]. In-frame knockout mutants were achieved by the replacement of target genes with a kanamycin resistance cassette. In the case of *A. baylyi*, deletion mutants were constructed as described previously [46]. Briefly, linear constructs of the kanamycin resistance cassette with 5’-overhangs homologous to the insertion site were amplified by PCR, where pKD4 was used as a template (see Supplemental Table S6 for primer details). Upstream and downstream regions homologous to *hisD* and *trpB* were amplified using primers with a 5’-extension complementary to the primers used to amplify the kanamycin resistance cassette. The three amplified products (upstream, downstream, and kanamycin) were combined by PCR, resulting in overhanging flanks with a kanamycin cassette. This PCR product was introduced into the *A. baylyi* WT strain. For this, the natural competence of *A. baylyi* was harnessed. The transformation was done by diluting 20 μl of a 16 h-grown culture in 1 ml lysogeny broth (LB). This diluted culture was mixed with 50 μl of the above PCR mix and further incubated at 30 °C with shaking at 200 rpm for 3 h. Lastly, 1 ml volume was pelleted, washed once with LB broth, plated on LB agar plates containing kanamycin (50 μg ml^−1^), and incubated at 30 °C for colonies to grow.

Conditional lethality of constructed auxotrophic mutations in MMAB medium was verified by inoculating 10^5^colony-forming units (CFU) ml^-1^of these strains into 1 ml MMAB medium with or without the focal amino acid (100 μM). After 24 h, their optical density (OD) was determined spectrophotometrically at 600 nm using FilterMax F5 multi-mode microplate reader (Molecular Devices) and the mutation was considered conditionally essential when growth did not exceed the OD_600nm_ of the uninoculated minimal medium [69, 71].

### Culture conditions and general procedures

A modified minimal media for *Azospirillum brasilense* (MMAB, [72]) was used for all experiments containing K_2_HPO_4_ (3 g L^−1^), NaH_2_PO_4_ (1 g L^−1^), KCl (0.15 g L^−1^), NH_4_Cl (1 g L^−1^), MgSO_4_ · 7H_2_O (0.3 g L^−1^), CaCl_2_ · 2H_2_O (0.01 g L^−1^), FeSO_4_ · 7H_2_O (0.0025 g L^−1^), Na_2_MoO_4_· 2H_2_O (0.05 g L^−1^), and 5 g L^−1^ D-glucose as a carbon source. 10 ml of trace salt solution was added per liter of MMAB media from the 1L stock. Trace salt stock solution consisted of filter-sterilised 84 mg L^−1^ of ZnSO_4_. 7H_2_O, 765 μl from 0.1 M stock of CuCl_2_. 2H_2_O, 8.1 μl from 1 M stock of MnCl_2_, 210 μl from 0.2 M stock of CoCl_2_. 6H_2_O, 1.6 ml from 0.1 M stock of H_3_BO_3_, 1 ml from 15 gL^−1^ stock of NiCl_2._

All strains were precultured in replicates by picking single colonies from LB agar plates, transferring them into 1 ml of liquid MMAB in 96-deep well plate (Eppendorf, Germany) incubating these cultures for 20 h. In all experiments, auxotrophs were precultured at 30 °C in MMAB, supplemented with 100 μM of the required amino acid. The next day, precultures were diluted to an optical density of 0.1 at 600 nm as determined by FilterMax F5 multi-mode microplate readers (Molecular Devices).

### Coculture experiment

Approximately 50 μl of preculture were inoculated into 1 ml MMAB, leading to a starting density of 0.005 OD. In the case of cocultures, donor and recipient were mixed in a 1:1 ratio by co-inoculating 25 μl of each diluted preculture without amino acid supplementation. Monocultures of both donors and recipient (with and without the focal amino acid) were inoculated using 50 μl of preculture. Cultures were incubated at a temperature of 30 °C and shaken at 220 rpm. Cell numbers were determined at 0 h and 24 h by serial dilution and plating. Donor strains were plated on MMAB agar plates, whereas recipients (auxotrophs) were differentiated on LB agar containing kanamycin (50 μg ml^−1^) to select for recipient strains. For key resources, see (Supplemental Table S7).

### Relative fitness measurement

To quantify the effect of amino acid cross-feeding on the fitness of the recipient, the number of colony-forming units (CFU) per ml was calculated for monoculture and coculture conditions at 0 h and 24 h. Each donor was individually paired with one of the recipients as well as grown in monoculture. Every combination was replicated four times. The relative fitness of each recipient was determined by dividing the growth of each genotype achieved in coculture by the value of its respective monoculture. Since different donor genotypes show inherent differences in growth, the growth of recipients in coculture was normalised to reduce to minimise potential effects of this variation. For this, growth of recipients in monoculture was first subtracted from its growth in coculture and then divided by the growth the respective donor genotype achieved in coculture.

### Amino acid supernatant experiment

To determine whether cross-feeding was mediated via compounds that have been released into the extracellular environment, the cell-free supernatants of donor genotypes were harvested and provided to receiver strains. To collect the supernatant, donors were grown in 2.5 ml MMAB in 48-deep well plates (Axygen, USA) and cultivated at 30 °C under shaking conditions (220 rpm). Supernatants were isolated in the mid-exponential growth phase and centrifuged for 10 min at 4,000 rpm. Then, supernatants were filter-sterilised (0.22 μm membrane filter, Pall Acroprep, USA) and stored at −20 °C. Meanwhile, receivers were grown in 1 ml MMAB in 96-well plates for 24 h. After adjusting the receiver OD_600nm_ to 0.1, 5 μl of the receiver culture was added to the replenished donor supernatant (total culturing volume: 200 μl, i.e. 160 μl donor supernatant + 40 μl MMAB) in 384-well plates (Greiner bio-one, Austria) (total: 50 μl culture). Four replicates of each comparison were grown for 24 h at 30 °C in a FilterMax F5 multi-mode microplate reader (Molecular Devices). MMAB without supernatant and monocultures of receiver strains were used as control. Growth was determined by measuring the optical density at 600 nm every 30 minutes, with 12 minutes of orbital shaking between measurements. OD_600nm_ was measured and analysed to calculate the maximum optical density (OD_max_) achieved by the receiver strain using the Softmax Pro 6 software (Table S8). For each donor supernatant-receiver pair, OD_max_ achieved by receivers with supernatant was subtracted from the values achieved by cultures grown without supernatant and normalised with the OD_600nm_, the respective donor strain had achieved at the time of supernatant extraction.

### Amino acid quantification by LC/MS/MS

All 20 proteinogenic amino acids in the culture supernatant were analysed. 100 μl of extracted supernatant was derivatised using the dansyl chloride method [73, 74]. Norleucine was added as an internal standard to the sample, and a calibration curve was generated by analysing all 20 amino acids at different concentrations. All samples were directly analysed via LC/MS/MS. Chromatography was performed on a Shimadzu HPLC system. Separation was achieved on an Accucore RP-MS 150 x 2.1, 2,6 μm column (Thermo Scientific, Germany). Formic acid 0.1% in 100% water and 80% acetonitrile were employed as mobile phases A and B. The mobile phase flow rate was 0.4 ml min^−1^, and the injection volume was 1 μl. Liquid chromatography was coupled to a triple-quadrupole mass spectrometer (ABSciex Q-trap 5500). Other parameters were: curtain gas: 40 psi, collision gas: high, ion spray voltage (IS): 2.5 keV, temperature: 550 °C, ion source gas: 1: 60 psi, ion source gas 2: 70 psi. Multiple reaction monitoring was used to determine the identity of the focal analyte. Analyst and Multiquant software (AB Sciex) were used to extract and analyse the data.

### Amino acid profile-based distance calculation using supernatant data

The similarity in the amino acid production profiles of different donor species was determined by calculating the Euclidean distance. If the amino acid production of a donor that is closely related to the focal auxotroph is given by *CR* = (*cr_1_,cr_2_,…,cr_20_*), and the amino acid production of a distantly related donor is given by *DR* = (*dr_1_,dr_2_,…,dr_20_*), the Euclidean distance between recipient and donor is:

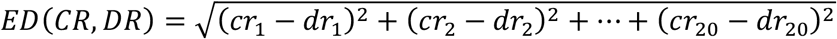

Index numbers (1-20) refer to individual amino acids.

### Phylogenetic tree construction and distance calculation

To cover a broad taxonomic diversity of donor strains, we chose 25 well-characterised species, belonging to four different phyla. The 16S rRNA gene sequences of 20 strains were retrieved from the NCBI GenBank and 5 strains from 16S rRNA sequencing (see supplementary method). The phylogenetic tree of this marker gene was generated using the maximum likelihood method in MEGA X software [75]. 16S rRNA gene locus sequences of all strains were aligned with MUSCLE. Maximum-likelihood (ML) trees were constructed using the Kimura 2-parameter model, where rates and patterns among mutated sites were kept at uniform rates, yielding the best fit. Bootstrapping was carried out with 1,000 replicates. The phylogenetic tree was edited using the iTOL online tool (Table S8) [76]. Pairwise phylogenetic distances between donor and receiver strains were extracted from a phylogenetic distance-based matrix. The resulting values quantify the evolutionary distance that separates the organisms.

### Reconstruction of metabolic networks

Genome-scale metabolic networks for all organisms (Table S5) were reconstructed based on their genomic sequences using the gapSeq software (version v0.9, https://github.com/jotech/gapseq) [77]. In brief, the reconstruction process is divided into two main steps. First, reactions and pathway predictions, and, second, gap-filling of the network to facilitate *in-silico* biomass production using flux balance analysis. For the reaction and pathway prediction step, all pathways from MetaCyc database [78] that are annotated for the taxonomic range of bacteria, were considered. Of each reaction within pathways, the protein sequences of the corresponding enzymes were retrieved from the SwissProt database [79] and aligned against the organism’s genome sequence by the TBLASTN algorithm [80]. An enzyme, and thus the corresponding reaction, was considered to be present in the organism’s metabolic network if the alignment’s bitscore was ≥ 200 and the query coverage ≥ 75%. Reactions were considered to be existing, if more than 75% of the remaining reactions within the pathway were predicted to be present by the BLAST-searches or if more than 66% of the key enzymes, which are defined for each pathway by MetaCyc, were predicted to be part of the network by the blast searches. As reaction database for model construction, we used the ModelSEED database for metabolic modeling [81].

The second step (i.e., the gap-filling algorithm of gapseq) solves several optimisation problems by utilising a minimum number of reactions from the ModelSEED database and adding them to the network to facilitate growth in a given growth medium. Here, the chemical composition of the M9 medium (which is qualitatively identical to MMAB) with glucose as sole carbon source was assumed.

### Calculating the genome-based metabolic distance of organisms

To estimate the pairwise metabolic distance between donor and recipient genotypes, the structure of their metabolic network was compared. For this, a flux balance analysis was performed on each individual metabolic network model with the biomass reaction flux as objective function. Subsequently, the biomass reaction flux was fixed to predicted maximum flux and a second flux balance analysis was performed to minimise the sum of absolute fluxes throughout the entire network [82]. Pairwise distances of flux distributions between organisms were calculated as the Euclidean distance between the predicted flux vectors. Only reactions with a non-zero flux in at least one of the two organisms were included in the distance approximations. In case a reaction was absent in one of the models, the flux was considered zero.

### In-silico *simulation of bacterial co-growth*

To further investigate the relationship between the metabolic distance between organisms and the likelihood of entering into a cross-feeding interaction, we extended our analysis to a larger number of bacterial organisms using *in-silico* co-growth simulations. For this, we reconstructed 818 bacterial metabolic network models as described above. The selected 818 organisms are the same as from the AGORA-collection, representing common members of the human gut microbiota [83]. For co-growth simulations, the models were merged in a pairwise manner, as described previously [84, 85]. The predicted flux values of the two individual biomass reactions (i.e. growth rate) were compared to the predicted growth rates of the respective models in monoculture, which enabled the prediction of potential growth benefits from metabolic interactions between both species. If at least one of two models was predicted to have a 20 % higher growth rate in coculture than in monoculture, the pair was considered as an interaction pair with growth advantage (Fig. 5). A logistic curve function of the form y = a/(1 + Exp[-b (x - c)]) was fitted to the data.

### Statistical data analysis

Normal distribution of data was evaluated employing the Kolmogorov-Smirnov test, and data was considered to be normally distributed when P > 0.05. Homogeneity of variance was determined using Levene’s test, and variances were considered homogenous if P > 0.05. Differences in the recipient growth in coculture versus monocultures were assessed with paired sample t-tests. P-values were corrected for multiple testing by applying the false discovery rate (FDR) procedure of Benjamini *et al.* [86, 87]. Linear regressions were used to assess the growth support of recipients in cocultures as a function of different variables (i.e. amino acid profile distance, phylogenetic distances, and metabolic distance). Spearman’s rank correlation was used to assess the relationship between amino acid production and growth of recipient as maximum density when cultured with donor supernatants. The relationship between each proxy tested and recipient growth was depicted as a 2D plane and analysed by fitting a linear regression. Regression analyses was also used to disentangle the effect of more than one interacting predictor variable. In these cases, the phylogenetic signal or amino acid produced was controlled for the respective other predictor variable (e.g. metabolic distance or amino acid production profile distance) used to predict the growth of recipient (Table S8).

## Acknowledgements

We thank the entire Kost lab (past and present) for useful discussion as well as Marita Hermann and Antje Moehlmeyer for technical assistance. We are grateful to Stefan Walter and Saskia Schuback (CellNanOS, MS facility) for help with quantifying the exo-metabolome, Heiko Vogel and Domenica Schnabelrauch (Department of Entomology, MPI-CE) for help with 16S rRNA sequencing, Ákos T. Kovács, (Technical University of Denmark; Denmark) for sharing two *Bacillus subtilis* strains and Michael Hensel, (Department of Microbiology, University of Osnabrück; Germany) for providing *Serratia ficaria*. This work was funded by the German Research Foundation (SPP1617, KO 3909/2-1: CK, SG), (SFB 944, P19: CK), (KO 3909/4-1: CK), DAAD GERLS program (GY, CK) and the University of Osnabrück (SG, LO, GY *EvoCell*: CK). CKa and SW acknowledge support by the German Research Foundation within the scope of the Excellence Cluster “Precision medicine in chronic inflammation” (EXC2167, sub-project RTF-VIII) and the Collaborative research center “Metaorganisms” (SFB1182, sub-project A1).

## Author contributions

SG and CKo conceived the project. SG, SS, and CKo designed the research. SG performed all experiments. SG and LO analysed the data. SG, LO, and CKo interpreted the data. SW calculated the genome-scale metabolic distance of tested strains and performed all *in-silico* analyses. SW and CKa carried out the gut microbiome *in-silico* data collection. GY isolated and phenotypically characterised the five environmentally-derived donor strains. SG and CKo wrote the manuscript, and all authors revised the manuscript. CKo provided resources and acquired funding.

