## Supplementary for "Metabolic dissimilarity determines the establishment of cross-feeding interactions in bacteria"

**Table of contents** **Page**

**Supplementary Figures**

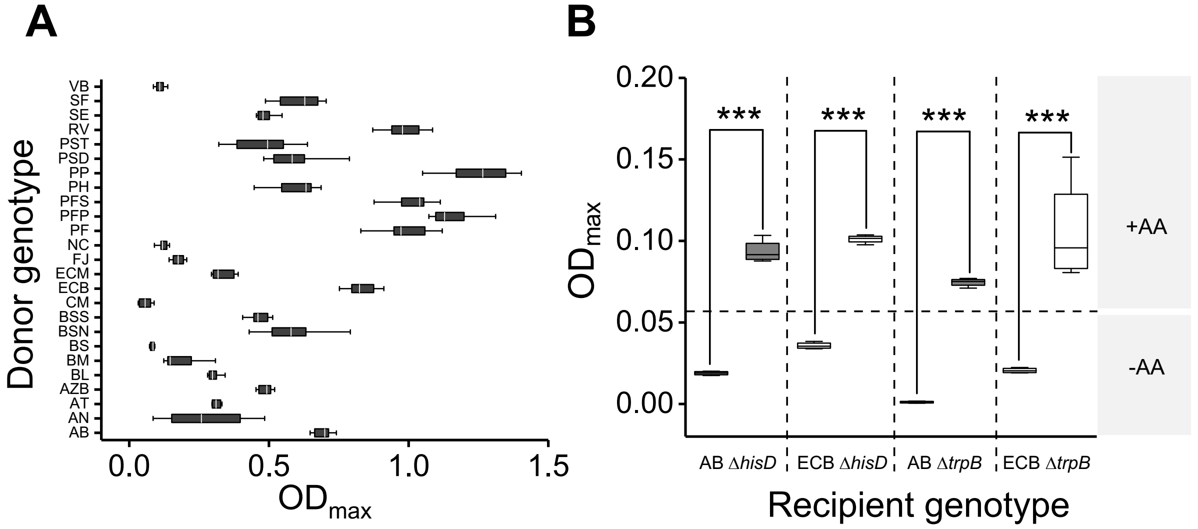

**Fig. S1. Donor and recipient growth after 24 h. (A)** Box plot shows growth of 25 donor strains in minimal medium (n=8). Donor genotypes were: *Acinetobacter baylyi* ADP1 (AB), *Arthrobacter nicotianae* (AN), *Agrobacterium tumefaciens* (AT), *Azospirillum brasilense* (AZB), *Bacillus licheniformis* (BL), *Bacillus megaterium* (BM), *Peribacillus simplex* (BS), *Bacillus subtilis* 3610 $comI^{Q12L}$ (BSN), *Bacillus subtilis* 168 $trpC^{+}$ (BSS), *Cupriavidus metallidurans* (CM), *Escherichia coli* BW25113 (ECB), *Escherichia coli* MG1655 (ECM), *Flavobacterium johnsoniae* (FJ), *Nocardia coeliaca* (NC), *Pseudomonas fluorescens* (PF), *Pseudomonas fluorescense* Pf-5 (PFP), *Pseudomonas fluorescens* SBW25 (PFS), *Pedobacter heparinus* (PH), *Pseudomonas putida* KT2440 (PP), *Pseudomonas syringae* DC 3000 (PSD), *Pseudomonas syringae* (PST), *Rahnella Victoriana* (RV), *Serratia entomophila* (SE), *Serratia ficaria* (SF), *Variovorax boronicumulans* (VB) (Table S5). **(B)** Growth of recipient strains (AB and ECB) auxotrophic for histidine (Δ*hisD)* or tryptophan (Δ*trpB)*, which have been cultivated in the presence (+AA) of the focal amino acid (100 µM) or without amino acid supplementation (-AA). Asterisks indicate the results of a paired sample t-test: *** P < 0.001, n=4. In both cases (A, B), growth was determined spectrophotometrically (OD 600 nm). Box plots: medians (lines within boxes), interquartile range (boxes), and 1.5x-interquartile range (whiskers).

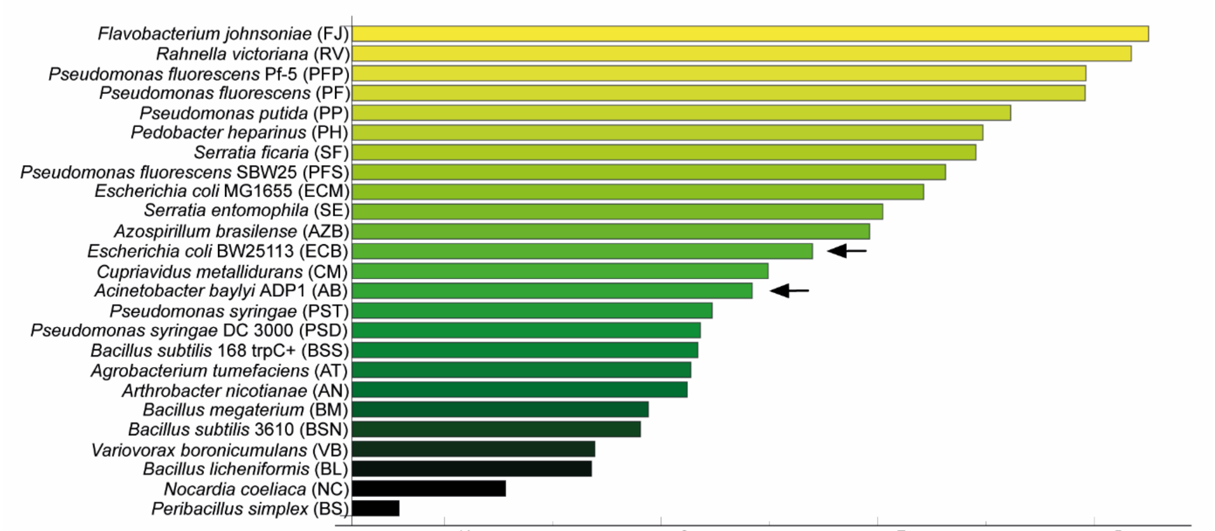

**Fig. S2. Ranking of donor strains according to their total amino acid production profile.** Amino acid production of 25 donor strains was quantified in mid-exponential growth and normalized by the number of colony forming units (CFUs). Bold names and arrows highlight the species used as recipients.

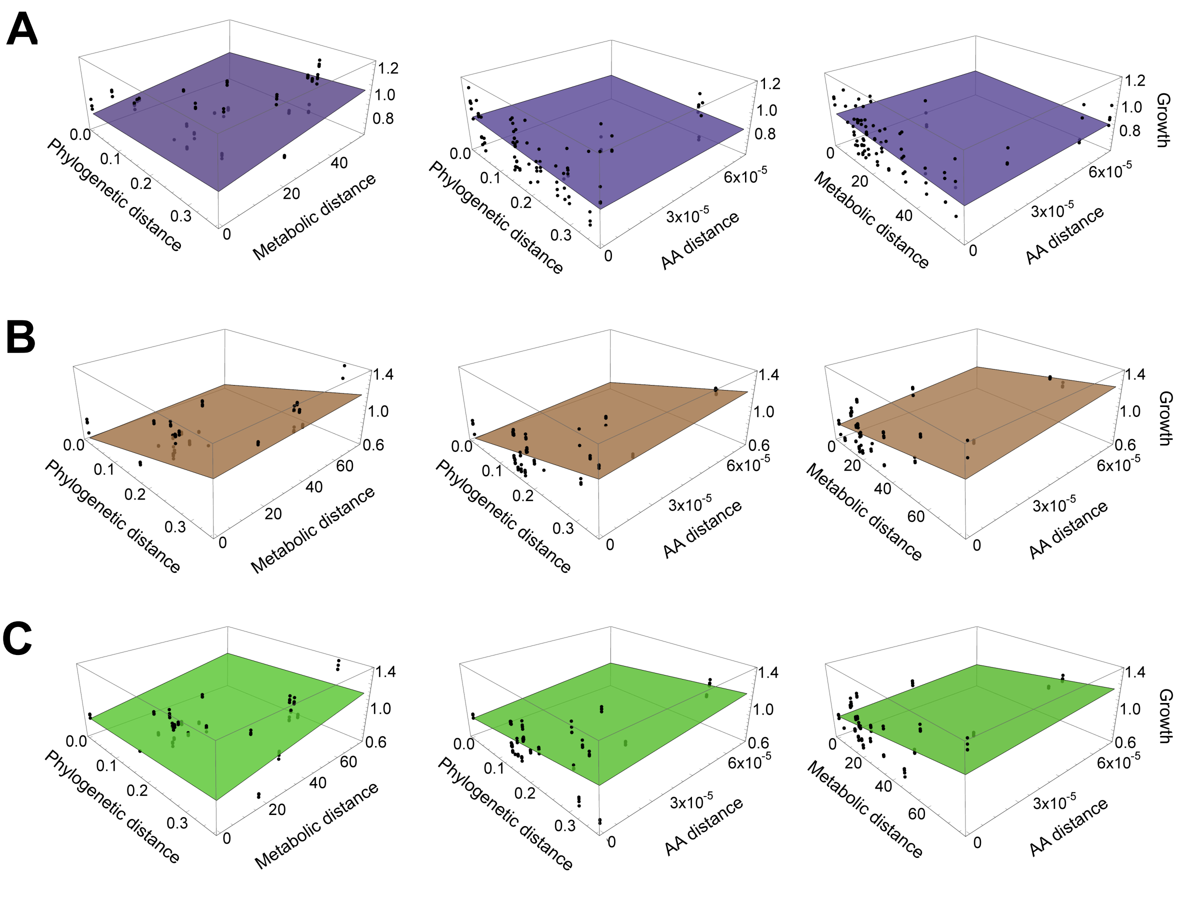

**Fig. S3. Multiple distance measures interactively explain recipient growth.** The plane depicts the linear regression between the growth of the **(A)** *E. coli* recipient auxotrophic for tryptophan (Δ*trpB*), **(B)** *A. baylyi* recipient auxotrophic for histidine (Δ*hisD*), and **(C)** *A. baylyi* recipient auxotrophic for tryptophan (Δ*trpB*) and the phylogenetic distance, the metabolic distance, and the amino acid profile distance between donor and recipient. Data points above the plane are shown in black.

### Supplementary Tables

**Table S1. Corrected regression analysis of the effect of amino acid profile distance (AAD), phylogenetic distance (PD), and metabolic distance (MD), on recipient growth.** To disentangle the effect of the individual parameters on the growth of auxotrophic recipients of *E. coli* (ECB) and *Acinetobacter baylyi* (AB), a linear regression for both MD and AAD was calculated with PD as the independent variable. The residuals (i.e., the variation not explained by PD) were then used as independent variables in the regressions. P and $R^2$ values represent significant regression coefficients. The results of the corrected regression remained significant for ECB recipients (Δ*hisD* and Δ*trpB)*, show marginally significant for AB (Δ*trpB*) but not in the case of AB (Δ*hisD*). Bold letters highlight significant effects (P ≤ 0.05).

| | | AAD corrected for PD | | MD corrected for PD | |
|---|---|---|---|---|---|
| Recipient | n | $R^2$ | P-value | $R^2$ | P-value |
| AB-his | 87 | 0.02 | 0.2 | 0.003 | 0.59 |
| AB-trp | 100 | 0.04 | 0.06 | 0.09 | **$3x10^{-3}$** |
| ECB-his | 80 | 0.17 | **0.013** | 0.08 | **0.01** |
| ECB-trp | 91 | 0.19 | **$1.4x10^{-5}$** | 0.07 | **0.011** |

**Table S2. Regression analysis between the amino acid profile distance (AAD), phylogenetic distance (PD), as well as metabolic distance (MD) and recipient growth, after correcting for the total amount of amino acids produced by donors (TAA).** To correct the effect that TAA would have on the ability of AAD, PD, and MD to predict growth of auxotrophic recipients, we first calculated a linear regression for both AAD, PD, and MD with TAA as the independent variable. The residuals (i.e., the variation not explained by TAA) were then used as independent variables in the regressions. Bold letters highlight significant effects (P ≤ 0.05).

| | | AAD corrected for TAA | | PD corrected for TAA | | MD corrected for TAA | |
|---|---|---|---|---|---|---|---|
| Recipient | n | $R^2$ | P-value | $R^2$ | P-value | $R^2$ | P-value |
| AB-his | 87 | 0.12 | **$9.3x10^{-4}$** | 0.37 | **$3.6x10^{-10}$** | 0.19 | **$2.1x10^{-5}$** |
| AB-trp | 100 | 0.2 | **$4.2x10^{-6}$** | 0.19 | **$5.6x10^{-6}$** | 1.25 | **$1.0x10^{-7}$** |
| ECB-his | 80 | 0.17 | **$1.4x10^{-4}$** | 0.04 | 0.06 | 0.1 | **0.0047** |
| ECB-trp | 91 | 0.24 | **$7.2x10^{-7}$** | 0.085 | **0.005** | 0.13 | **0.00042** |

**Table S3. Regression analysis between the amino acid profile distance (AAD), phylogenetic distance (PD), and metabolic distance (MD) on recipient growth, after correcting for the amount of the focal amino acid donors produced (FAA).** To correct the effect the FAA would have on the ability of AAD, PD, and MD to explain the growth of auxotrophic recipients, we first calculated a linear regression for both AAD, PD, and MD with FAA as the independent variable. The residuals (i.e., the variation not explained by FAA) were then used as independent variables in the regressions. Bold letters highlight significant effects ($P \leq 0.05$).

| | | AAD corrected for FAA | | PD corrected for FAA | | MD corrected for FAA | |
|---|---|---|---|---|---|---|---|
| Recipient | n | $R^2$ | P-value | $R^2$ | P-value | $R^2$ | P-value |
| AB-his | 87 | 0.039 | 0.065 | 0.33 | **$6.8x10^{-9}$** | 0.16 | **$8.5x10^{-5}$** |
| AB-trp | 100 | 0.05 | **0.02** | 0.08 | **$4.3x10^{-3}$** | 0.15 | **$5.8x10^{-5}$** |
| ECB-his | 80 | 0.06 | **0.02** | 0.045 | 0.059 | 0.059 | **0.03** |
| ECB-trp | 91 | 0.11 | **$1.5x10^{-3}$** | 0.06 | **0.02** | 0.06 | **0.02** |

**Table S4.** Logistic equation (y = a/(1 + Exp[-b (x - c)])) parameter estimation and ANOVA table for *in-silico* flux-balance-analysis. The result is confirmed formally by noting a logistic equation in which estimation of each parameter, standard error, t-statistic and P-values per parameter provides test values. ANOVA was conducted using the same data set for degree of freedom (DF), Sum-of-squares (SS) and mean squares (MS) to check the data distribution as in the following table (related to Figure 5).

**Logistic equation parameter estimation table**

**For phylogenetic distance**

| | **Estimate** | **Standard error** | **t-statistic** | **P-value** |
|---|---|---|---|---|
| a | 0.544782 | 0.005468 | 99.6289 | $<10^{-4}$ |
| b | 20.0741 | 1.56481 | 12.8285 | $<10^{-4}$ |
| c | 0.08729 | 0.0038703 | 22.5548 | $<10^{-4}$ |

**For metabolic distance**

| | **Estimate** | **Standard error** | **t-statistic** | **P-value** |
|---|---|---|---|---|
| a | 0.494469 | 0.003122 | 158.382 | $<10^{-4}$ |
| b | 0.9018 | 0.0826011 | 10.9176 | $<10^{-4}$ |
| c | 3.77157 | 0.0996954 | 37.8309 | $<10^{-4}$ |

**ANOVA table**

**For phylogenetic distance**

| | **DF** | **SS** | **MS** |
|---|---|---|---|
| Model | 3 | 127.346 | 42.4488 |
| Error | 535 | 5.08054 | 0.00949634 |
| Uncorrected total | 538 | 132.427 | |
| Corrected total | 537 | 14.0497 | |

**For metabolic distance**

| | **DF** | **SS** | **MS** |
|---|---|---|---|
| Model | 3 | 200.626 | 66.8754 |
| Error | 885 | 6.43082 | 0.00726646 |
| Uncorrected total | 888 | 207.057 | |
| Corrected total | 887 | 11.6224 | |

**Table S5.** Overview of bacterial strains used in the study.

| Strains | Source | Identifier |
|---|---|---|
| **Donors** | | |
| *Acinetobacter baylyi* ADP1 (AB) | [1] | [2] |
| *Arthrobacter nicotianae* (AN) | German Collection of Microorganisms and Cell Cultures, DSMZ | DSM 20123 |
| *Agrobacterium tumefaciens* (AT) | Lab stock | N/A |
| *Azospirillum brasilense* (AZB) | German Collection of Microorganisms and Cell Cultures, DSMZ | DSM1690 |
| *Bacillus licheniformis* (BL) | This study | Soil isolate from sample site coordinates 50.906557, 11.505631 |
| *Bacillus megaterium* (BM) | German Collection of Microorganisms and Cell Cultures, DSMZ | DSM 32 |
| *Peribacillus simplex* (BS) | This study | Soil isolate from sample site coordinates 50.906557, 11.505631 |
| *Bacillus subtilis* 3610 *comI*$^{Q12L}$ (BSN) | [3] | Provided by Ákos T. Kovács, DTU |
| *Bacillus subtilis* 168 *trpC*$^{+}$ (BSS) | [4] | Provided by Ákos T. Kovács, DTU |
| *Cupriavidus metallidurans* (CM) | This study | Soil isolate from sample site coordinates 50.906557, 11.505631 |
| *Escherichia coli* BW25113 (ECB) | [5] | *E. coli* Genetic resources at Yale CGSC, The Coli Genetic Stock Center |
| *Escherichia coli* MG1655 (ECM) | German Collection of Microorganisms and Cell Cultures, DSMZ | DSM 18039 |
| *Flavobacterium johnsoniae* (FJ) | German Collection of Microorganisms and Cell Cultures, DSMZ | DSM 2064 |
| *Nocardia coeliaca* (NC) | This study | Soil isolate from sample site coordinates 50.906557, 11.505631 |
| *Pseudomonas fluorescens* (PF) | German Collection of Microorganisms and Cell Cultures, DSMZ | DSM 289 |
| *Pseudomonas fluorescense* Pf-5 (PFP) | Lab stock | N/A |
| *Pseudomonas fluorescens* SBW25 (PFS) | Lab stock | [6] |
| *Pedobacter heparinus* (PH) | German Collection of Microorganisms and Cell Cultures, DSMZ | DSM 2366 |
| *Pseudomonas putida* KT2440 (PP) | Lab stock | DSM 6125 |
| *Pseudomonas syringae* pv. tomato DC 3000 (PSD) | Lab stock | N/A |
| *Pseudomonas syringae* subsp. syringae van Hall 1902 (PST) | German Collection of Microorganisms and Cell Cultures, DSMZ | DSM 50315 |

| | | |
|---|---|---|
| *Rahnella victoriana* (RV) | German Collection of Microorganisms and Cell Cultures, DSMZ | DSM 27397 |
| *Serratia entomophila* (SE) | German Collection of Microorganisms and Cell Cultures, DSMZ | DSM 12358 |
| *Serratia ficaria* (SF) | Lab stock | Provided by Department of Microbiology, University of Osnabrück |
| *Variovorax boronicumulans* (VB) | This study | Soil isolate from sample site coordinates 50.906557, 11.505631 |
| **Recipients** | | |
| *Acinetobacter baylyi* ADP1, *ΔhisD::kanR* | This study | N/A |
| *Acinetobacter baylyi* ADP1, *ΔtrpB::kanR* | This study | N/A |
| *Escherichia coli* BW25113, *ΔhisD::kanR* | This study | N/A |
| *Escherichia coli* BW25113, *ΔtrpB::kanR* | This study | N/A |

**Table S6. Primers used for the construction of *Acinetobacter baylyi* auxotrophs and for the identification of isolated strains.** To create auxotrophic genotypes, biosynthetic genes were replaced with a kanamycin resistance cassette. Strains were identified by sequencing using 16S rRNA. UF = upstream forward, UR = upstream reverse, DF = downstream forward, DR = downstream reverse, FP = forward primer, RP = reverse primer.

| Gene | Primer | Sequence (5'-3') |
|---|---|---|
| **Kanamycin resistance cassette** | UF | TGTAGGCTGGAGCTGCTTC |
| | UR | CATATGAATATCCTCCTTA |
| ***hisD*** | UF | TATGCAAGCCTTGGTGAGCA |
| | UR | GAAGCAGCTCCAGCCTACACAGCCTCTTCCACTTGA |
| | DF | AAGGAGGATATTCATATGGTAACTGCTCTACGGGG |
| | DR | ATGCGTCTGCCTGATCTACC |
| ***trpB*** | UF | AACCACACACGCTTTTGCAG |
| | UR | GAAGCAGCTCCAGCCTACAGCTGATCCACATTGGACT |
| | DF | TAAGGAGGATATTCATATGACGTGATGTGGAAATGG |
| | DR | AGTTGGGGCTGGATGTCTTG |
| **Eubacterial 16S rRNA primers** | FP | AGAGTTTGATCCTGGCTCAG |
| | RP | ACGGCTACCTTGTTACGACTT |

**Table S7.** Overview of reagents or material used in the study.

| Reagent or resource | Source | Identifier |
|---|---|---|
| Lysogeny broth (LB), Lennox | Carl Roth GmbH | Catalog # X964.1 |
| Agar-Agar Kobe | Carl Roth GmbH | Catalog # 5210.2 |
| Dipotassium hydrogen phosphate | Carl Roth GmbH | Catalog # 26931.263 |
| Sodium dihydrogen phosphate | Carl Roth GmbH | Catalog # T879.2 |
| Magnesium sulphate heptahydrate | Carl Roth GmbH | Catalog # P027.2 |
| Potassium chloride | VWR | Catalog # 26764.260 |
| Calcium chloride dihydrate | Carl Roth GmbH | Catalog # 5239.2 |
| Ammonium chloride | VWR | Catalog # 21236.267 |
| Iron (II) sulphate heptahydrate | Merck | Catalog # 3965 |
| Manganese chloride | AppliChem | Catalog # A2087.0100 |
| Cobalt chloride hexahydrate | AppliChem | Catalog # A2087.0100 |
| Boric acid | Carl Roth GmbH | Catalog # 6943.3 |
| Nickel chloride | AppliChem | Catalog # A3917.0100 |
| Zinc sulphate heptahydrate | Carl Roth GmbH | Catalog # K301.1 |
| Copper chloride dihydrate | AppliChem | Catalog # 131264.1210 |
| Glucose | Carl Roth GmbH | Catalog # 6887.1 |
| Kanamycin | Carl Roth GmbH | Catalog # T832.2 |
| Sodium carbonate | Merck | Catalog # 1063920.500 |
| Acetonitrile | Sigma-Aldrich | Catalog # 271004 |
| Formic acid | Acros-Organics | Catalog # 270480250 |
| L-Alanine | AppliChem reagents | Catalog # A3690,0100 |
| L-Arginine monohydrochloride | AppliChem reagents | Catalog # A3709,0100 |
| L-Asparagine monohydrochloride | AppliChem reagents | Catalog # A3721,0100 |
| L-Aspartic acid | AppliChem reagents | Catalog # A3715,0250 |
| L-Cysteine monohydrochloride | AppliChem reagents | Catalog # A3694,0050 |
| L-Glutamine monohydrochloride | AppliChem reagents | Catalog # A3734,0100 |
| L-Glutamic acid monohydrochloride | AppliChem reagents | Catalog # A3712,0250 |
| Glycine | AppliChem reagents | Catalog # A1067,1000 |
| L-Histidine monohydrochloride | AppliChem reagents | Catalog # A3733,0100 |
| L-Isoleucine | AppliChem reagents | Catalog # A1440,1000 |
| L-Leucine | AppliChem reagents | Catalog # A3460,0100 |
| L-Lysine monohydrochloride | AppliChem reagents | Catalog # A3466,0100 |
| L-Methionine | AppliChem reagents | Catalog # A1340,0100 |
| L-Phenylalanine | AppliChem reagents | Catalog # A3442,0100 |
| L-Proline | AppliChem reagents | Catalog # A3453,0100 |
| L-Serine | AppliChem reagents | Catalog # A3973,0100 |
| L-Threonine | AppliChem reagents | Catalog # A3969,0100 |
| L-Tryptophan | AppliChem reagents | Catalog # A3445,0100 |
| L-Tyrosine disodium salt hydrate | Sigma | Catalog # T1145-25G |
| L-Valine | AppliChem reagents | Catalog # A1637,1000 |
| 48-well deep well plates | Axygen | Catalog # P-5ML-48-C-S |
| 96-well deep well plates | Eppendorf | Catalog # 0030506308D |
| 384-well plates | Greiner bio-one | Catalog # 781185 |
| Plate sealer | Greiner bio-one | Catalog # 676070 |
| Petri dish | Greiner bio-one | Catalog # 633180 |
| 96 well Filter plates (0.2 µm PTFE membrane) | Pall AcroPrep | Catalog # 8047 |

**Table S8.** List of software used in the study.

| Software | Source | Identifier |
|---|---|---|
| Origin Pro 2017 | OriginLab, Northampton, MA | https://www.originlab.com/index.aspx?go=Products/Origin |
| IBM SPSS statistics 26 | IBM Corp. Released 2019. IBM SPSS Statistics, Version 26.0. Armonk, NY: IBM Corp | https://www.ibm.com/support/pages/downloading-ibm-spss-statistics-26 |
| MEGA X | [7] | https://www.megasoftware.net/ |
| iTOL | [8] | https://itol.embl.de/ |
| Softmax Pro 6 software | Molecular Devices | N/A |
| R version 3.5.3 | [9] | http://www.R-project.org |
| Mathematica | N/A | https://www.wolfram.com/mathematica/ |

### Supplementary Methods

#### Soil sampling, bacterial isolation, 16S rRNA amplification, and taxonomic affiliation

Soil samples were collected from a meadow site in Jena, Germany. A soil column of 50 mm diameter and 50 mm length was collected using the soil sampler and transported to the lab without disturbing its structure. To process the soil sample, the structure was carefully intersected and three 1 mg soil particles, that were spaced apart by 2.5 cm, were picked for bacterial strain isolation. Soil particles were then suspended in 500 µl of saline solution (0.9% NaCl) and shaken for 30 mins. After that, 100 µl were transferred into 400 µl saline solution and vortexed vigorously for 5 min. Next, 100 µl of the previously prepared dilutions were spread on agar plates (in three replicates). The medium used for isolation has a MMAB composition, with fructose as a sole carbon source, and supplemented with; twenty amino acids (200 µM), five different vitamins (i.e. riboflavin ($B_2$), cobalamin ($B_{12}$), biotin ($B_7$), thiamine ($B_1$), and pyridoxine ($B_6$), each at 1 µM). In addition, 100 µg $ml^{-1}$ of the fungicide Nystatin was added to the media. Plates were incubated for three days at room temperature. After incubation, all colonies were picked and purified three times on agar plates to finally isolate a single colony. The resulting isolates were inoculated into 96-well plates containing the previously used medium and incubated under shaking conditions for 30 hours. Finally, glycerol was added to a final concentration of 50% (vol/vol) to prepare stocks that were frozen at -80º C until further use.

To characterize the five bacterial isolates (i.e. *Peribacillus simplex, Variovorax boronicumulans, Cupriavidus metallidurans, Bacillus licheniformis* and *Nocardia coeliaca*), the focal strains were revived to PCR-amplify and sequence part of the housekeeping 16S rDNA gene. For this, 1 µl of bacterial biomass from an overnight-grown culture was directly used as PCR template. The 16S rRNA gene was amplified using the general eubacterial primers FP and RP [10], (Table S6) on a Biometra TProfessional Thermocycler (Biometra, Jena, Germany) in 20 µl reaction volumes using JumpStart™ REDTaq ReadyMix (Sigma Aldrich). 96-well plates were sealed with adhesive film. The following parameters were used: 3 min at 94 °C, followed by 32 cycles of 40 s at 94 °C, 60 s at 65 °C and 60 s at 72 °C, and a final extension step of 4 min at 72 °C. PCR products were checked on an agarose gel, purified, treated with Shrimp alkaline phosphatase (Illustra, Fischer scientific), and sequenced using an ABI 3730XL capillary DNA sequencer (Applied Biosystems, USA) in the Department of Entomology, Max-Planck-Institute for Chemical Ecology in Jena, Germany.

The forward and reverse 16S rRNA gene sequences were assembled and manually curated using Microsoft Word Office 2011 to generate contigs after trimming the poor sequences at both ends. Similarity-based searches were carried out using NCBI BLASTn [11] for taxonomic assignment of 16s rRNA sequences. The species of the five focal strains have been taxonomically identified based on sequence similarity to their nearest neighbor by using minimum query coverage of 98% and minimum identity values of 99%.

The 16S rRNA gene sequences of the 5 soil-derived strains used in this study have been deposited in Genbank under the following accession number, *Peribacillus simplex* (MW073531), *Variovorax boronicumulans* (MW073532), *Cupriavidus metallidurans* (MW073533), *Bacillus licheniformis* (MW073534), and *Nocardia coeliaca* (MW080346-MW080347).
